# High-frequency changes in single-trial visual evoked potentials for unattended stimuli in chronic schizophrenia

**DOI:** 10.1101/2021.11.09.467985

**Authors:** Lech Kipiński, Andrzej Maciejowski, Krzysztof Małyszczak, Witold Pilecki

## Abstract

**Background:** Patients with schizophrenia reveal changes in information processing associated with external stimuli, which is reflected in the measurements of brain evoked potentials. We discuss actual knowledge on electro- (EEG) and magnetoencephalographic (MEG) changes in schizophrenia.

**New method:** The commonly used averaging technique entails the loss of information regarding the generation of evoked responses. We propose a methodology to describe single-trial (non-averaged) visual evoked potentials (VEP) using spectral and statistical analyses. We analysed EEG data registered in the O1-Cz and O2-Cz leads during unattended pattern-reversal stimulation, collected from a group of adult patients with chronic schizophrenia, and compared them to those of healthy individuals. Short-time single-trial VEP were transformed to the frequency domain using the FFT algorithm. Changes of the spectral power were visualized using spectrograms which were created by stacking single-trial spectra across all trials. Measures of the absolute and the relative spectral power were calculated and compared statistically.

**Results:** In schizophrenia, the energy density of VEP oscillations is shifted towards higher (gamma) frequencies, compared to healthy individuals. These differences are statistically significant in all analysed frequency bands for the relative power. This indicates distorted early processing of visual stimuli in schizophrenia.

**Comparison with existing methods:** The main advantage of the presented methodology is its simplicity and ease of interpretation of obtained results. The presented observations complement the knowledge on gamma oscillations acquired from computationally more complex methods of time–frequency analysis.

**Conclusions:** High-frequency changes for single-trial VEPs are detected in chronic schizophrenia.

## 1. Introduction

Schizophrenia exhibits a number of disorders of higher nervous functions and it is believed that the abnormalities in the synthesis of the perceived sensory impressions may underlie mental disorders in patients with schizophrenia. Therefore, plenty of research have been devoted to the search for biological markers of schizophrenia among the parameters obtained from electro-(EEG) and magnetoencephalographic (MEG) measurements and reflecting the response of the human brain to external stimuli. These efforts have been going on since the early 1980s (Connolly et al., 1983) and continue to this day (van der Stelt and Belger, 2007; Kim et al., 2020).

### 1.1. Resting-state EEG/MEG

Even the spontaneous electroencephalograms demonstrate abnormalities in schizophrenia (Narayanan et al., 2014; Renaldi et al., 2019). Such phenomena can be described based on the spectral EEG/MEG analysis, which quantifies the traditional delta, theta, alpha, beta or gamma waves by the use of absolute or relative spectral power measures in selected frequency bands. Spectral abnormalities in the EEG measurements as a diagnostic marker for schizophrenia were widely reviewed by Boutros et al. (2008). In that work, authors concluded that most of the research reports the increased preponderance of slow EEG rhythms in schizophrenia patients and this effect remained consistent in unmedicated patients. In turn, in a recent review (Newson and Thiagarajan, 2019) authors concluded that the most dominant pattern of changing schizophrenia is spectral power increases across lower frequencies (delta and theta) and decreases across higher frequencies (alpha, beta, and gamma). Slowing of the EEG would reflect an impaired subcortical synchronization system including the mesencephalic reticular formation, nucleus reticularis and the thalamus (Boutros et al., 2008; Kirino, 2004). The increased delta absolute power is observed from the beginning of psychotic disorders and its spatial changes during treatment may have a positive predictive value (Renaldi et al., 2019). Unfortunately, based on spontaneous electromagnetic brain activity, the characteristic patterns of power change within specific frequency bands are not necessarily unique to schizophrenia or any psychiatric disorder but show substantial overlap across disorders as well as variability within disorders (Newson and Thiagarajan, 2019). For example, an aberrant low-frequency activity (as well as changes in theta, and slow and fast alpha activity) is observable in other psychotic disorders, such as bipolar disorder (Narayanan et al., 2014).

### 1.2. Evoked activity

In many aspects, more interesting information is provided by the registration of the brain bioelectromagnetic activity related to the processing of external stimuli. Just as the spontaneous EEG or MEG reflects the resting state, processing stimuli forces changes in the functioning of the neural network related to their perception, synthesis and classification (or not) of the information it contains. These changes are described by other EEG/MEG-derived measures, such as event-related responses (ERR), i.e. event-related potentials (ERP) in EEG or event-related fields (ERF) in MEG and evoked oscillations. Based on them, neurophysiological deficits in both auditory and visual perception are observable from the very early phase of sensory processing (P50, N100 waveforms) to the relatively late phase (P300, N400 components) in patients with schizophrenia (Onitsuka et al., 2013). Moreover, the neural synchrony measured by auditory steady-state responses (ASSR) and steady-state visual evoked potentials (SSVEP) shows several anomalies in schizophrenia patients (Brenner et al., 2009).

#### 1.2.1. Cognitive event-related responses

Some of pathophysiological theories of schizophrenia highlight the role of altered brain connectivity manifesting functionally through an aberrant control of synaptic plasticity (Stephan et al., 2006). This results in malfunctioning of neural circuits, preventing proper cognitive integration, which appears to be the basis for this disorder (Spencer et al., 2003; Onitsuka et al., 2013). Therefore, studying cognitive ERPs to measure this dysconnectivity is of great importance for understanding the pathophysiology of schizophrenia. The best known cognitive event-related component is the P300 (P3) waveform, and its great role as a diagnostic marker, prognostic indicator or endophenotype in schizophrenia is described in many papers— for a systematic review see Galderisi et al. (2009). The main finding is the amplitude reduction of the P300 wave, obtained in the oddball procedure, which is connected to attention-mediated sensory processing deficits (Oribe et al., 2013; Hamilton et al., 2018). Unfortunately, this phenomenon is well-replicated but completely non-specific for schizophrenia (Ergen et al., 2008). Another cognitive ERP, designed to access functioning of the sensory memory system is the mismatch negativity (MMN) potential and its deficits in schizophrenia have been reported, for example, by Javitt et al. (2001); Kim et al. (2020). Such deficits occur in the absence of directed attention. They reflect working memory dysfunction and are associated with poor functioning of schizophrenic patients (Light and Braff, 2005; Hamilton et al., 2018), yet, they, too, are non-specific (Erickson et al., 2015). In the work of Xia et al. (2020), association of cognitive and P50 suppression deficits in chronic schizophrenia is shown. The MMN component and the P50 wave are considered as potential endophenotypes in schizophrenia (Kim et al., 2020).

#### 1.2.2. Early-stage potentials

In contrast to the research on deficits in higher cognitive functions (including attention, executive function and memory) in schizophrenia some authors emphasize that changes in the basic perceptual processes in schizophrenia may represent a core impairment (Javitt et al., 2001; Uhlhaas and Mishara, 2007). It is clearly visible in visual evoked potentials (VEP) as a reduced amplitude of the P100 wave, which has repeatedly been demonstrated in schizophrenia (Foxe et al., 2001; Yeap et al., 2006; Martinez et al., 2012; González-Hernández et al., 2015). These changes over the parieto-occipital scalp suggest that schizophrenia is associated with impairment of early dorsal visual stream processing (Foxe et al., 2001; Lalor et al., 2012; Oribe et al., 2013). Early visual processing deficits in schizophrenia are also described in the pattern-reversal paradigm as an atypical VEP waveform morphology (Knebel et al., 2011; Babhulkar et al., 2017) or elongated latency of the P100 (Małyszczak et al., 2003; Rady et al., 2011). For the face-recognition paradigm both the P100 and N170 waves reveal decrements in their amplitudes, which suggests global deficits in early visual processing that might underlie the decreased amplitudes and higher onset latencies of the later P300 and N400 components (Campanella et al., 2006). These perceptual deficits are evident in chronic schizophrenia and the most recent studies show that they are associated with social cognitive performance in recent-onset schizophrenia (McCleery et al., 2020). A systematic review on early-stage visual processing in schizophrenia is given in Butler and Javitt (2005).

#### 1.2.3. Neural oscillations

Classical theories of sensory processing view the brain as a passive, stimulus-driven device. The event-related brain activity is considered as a stimulus-determined signal hidden in the ongoing EEG/MEG background. Newer theories, in contrast, view perception as an active, selective process controlled by top-down influences that include attentional processes, working memory and behavioural context (Engel et al., 2001). Thus, EEG/MEG activity is considered as generated by interactions between higher and lower cortical areas, described by neural oscillations which influence perception, enhancing some processes and suppressing others. In this concept, EEG/MEG fluctuations are not treated as background noise, but as ongoing signals with stimulus-driven phase and amplitude. Then, the perceptual processing can be understood as reorganization of the ongoing oscillatory activity, which manifests as an event-related desynchronization (ERD) or event-related synchronization (ERS) (Başar et al., 1999; Pfurtscheller, 1992; Pfurtscheller and Lopes da Silva, 1999; Żygierewicz et al., 2008). The relationship between cortical oscillations and schizophrenia is described widely and it is well proved that abnormalities in the synchronized oscillatory neural activity play a central role in the pathophysiology of schizophrenia (Spencer et al., 2003; Uhlhaas et al., 2010; Uhlhaas and Singer, 2010; van der Stelt and Belger, 2007).

Although cognitive functions are associated with synchronized oscillatory activity in the theta-, alpha, beta-, and gamma-band, suggesting a functional mechanism of neural oscillations in cortical networks, the best known role in sensory and cognitive processing, cortico-cortical transmission, and integration of information across neural networks is their association with the gamma oscillations (Light et al., 2006; Fries et al., 2007; Uhlhaas and Singer, 2010). This 30–100-Hz activity plays an important role in neuronal communication and synaptic plasticity, and is associated with attention and memory (Buzsáaki and Draguhn, 2004; Jensen et al., 2007). For example, attention can enhance synchrony or gamma-band oscillations in neurons representing the attended stimulus (Engel et al., 2001), and the gamma-band synchronization in visual cortex predicts the speed of change detection (Womelsdorf et al., 2006). The importance of gamma-synchronization for the impaired brain integration as a pathophysiological feature of schizophrenia is widely described in the literature (see, e.g., Lee et al., 2003; van der Stelt et al., 2004; Spencer et al., 2008). Recent MEG studies show that individuals with psychotic disorder prone to visual hallucinations will exhibit deviant functions within early visual cortex, and that aberrant contextual influences on visual perception will involve higher-level visual cortical regions and are associated with visual hallucinations (Klein et al., 2020). This suggests existing of perceptual mechanisms of visual hallucinations and illusions in schizophrenia. Moreover, in visual-priming experiment in chronic schizophrenia, impaired high-frequency (gamma) oscillations and event-related magnetic fields in thalamo-occipital cortices were found (Sauer et al., 2020).

Interestingly, high gamma power is phase-locked to theta oscillations in human neocortex (Canolty et al., 2006). In turn, for unattended perception, posterior alpha oscillations provide a mechanism for prioritizing and ordering visual input, and gamma oscillations phase-locked to them keep competing unattended representations apart in time, thus creating a sequence of perceptual cycles (Jensen et al., 2012). In addition, the high-frequency activity in human visual cortex is modulated by visual motion strength (Siegel et al., 2007). The utility of stimulus-related electroencephalographic measures in schizophrenia studies, such as event related potentials and evoked gamma oscillations, has been reviewed by van der Stelt and Belger (2007).

### 1.3. Methods of EEG/MEG signal processing

At the end of our introductory considerations, some emphasis should be placed on the methodology used to determine physiological measures from the recorded EEG/MEG signals.

Without system-changing events, like the external stimuli which activate some pathways in brain functional network, encephalograms can be considered as stationary signals (B. A. Cohen, 1977). Then, the simplest quantifiable electrophysiological measures for spontaneous EEG/MEG is offered by the spectral analysis. The most useful method to obtain the representation of a brain signal in the frequency domain is well-known fast Fourier transform (FFT) algorithm (Burrus, 2012). In this way such spectral measures like the absolute spectral power and the relative spectral power can be calculated. Due to its simplicity and relatively low computational cost it is widely used in neuroscience and its utility in schizophrenia research is well documented (see, e.g., Fenton et al., 1980; Newson and Thiagarajan, 2019).

In turn, ERPs/ERFs are derived by arithmetically averaging multiple registrations with respect to the onset of a repeated stimulation. Such averaging is simple and widely applied to extract the useful signal embedded in the background noise (the latter including spontaneous brain activity) but it is based on the assumption that the evoked response is deterministic and the ongoing noise is random and normally distributed (de Munck et al., 2004; Truccolo et al., 2002) with zero expected value. Unfortunately, this method does not take into account changes in the brain functional state that take place during the examination and that come from perceiving new stimuli and recognizing them, getting used to them, from perceiving other stimuli and from some perceptual changes during examination (Sielużycki and Kordowski, 2014; König et al., 2015). Although such information is being lost during averaging, we can obtain it by analysing the singletrial responses, which is more difficult and requires more advanced algorithms (Kipiński and Maciejowski, 2010).

Encephalographic signals are nonstationary, i.e., their properties change over time (Kaplan et al., 2005; Kipiński et al., 2011; Kipiński, 2011). They can be evaluated by the time–frequency analysis methods, with the use of tools such as the short-time (windowed) Fourier transform (STFT), the Wigner–Ville distribution (WVD), the Hilbert transform (HT), the wavelet transforms (WT) or the matching pursuit (MP) decomposition (Başar et al., 2001; Sielużycki et al., 2009; Jörn et al., 2011; Wacker and Witte, 2013; Baumgartner et al., 2013). Other advanced techniques dedicated to EEG/MEG signal description are based on fractal or entropy analysis (Racz et al., 2020; Trujillo, 2019), the application of independent component analysis (ICA) and similar techniques (Jung et al., 2001; Anemüller et al., 2003; Hsu et al., 2018) or various statistical methods (Georgiadis et al., 2005; Sielużycki and Kordowski, 2014; Galka et al., 2011; Matsuda and Komaki, 2017a,b). Some of them use both time-varying spectrum approximation and statistical time series modelling (Kipiński, 2007; Kipiński and Kordecki, 2021).

In schizophrenia research, wavelet transforms have been used successfully. In line with the theory of oscillation, description of nonlinear brain dynamics is possible via the time–frequency analysis, allowing for the monitoring of time changes in oscillations energy (Başar et al., 1999). Such analysis is crucial to the description of the functional-complexity of stimuli-induced neural networks. Wavelets offer the possibility of the single-trial EEG/MEG signal decomposition and computation of event-related spectral measures like phase locking factor or evoked power. This methodology has been applied in many schizophrenia studies (see, e.g., Ergen et al., 2008; Spencer et al., 2008; Donkers et al., 2013; Yaesoubi et al., 2017; Roach et al., 2019, and references therein).

Of course, time–frequency methods have some limitations. Most of them are related to the time–frequency resolution, which has consequences in building an adequate representation of the signal in the time–frequency domain based on these methods. In STFT, it is difficult to correctly assume the optimal window width for an unknown signal. For the quadratic transforms like WVD, one needs to minimize cross-terms effects. In wavelet representations, the trade-off between the time resolution and the frequency resolution is emphasized by the intrinsic scaling of the mother wavelet. Besides, one must also deal with the choice of the optimal wavelet shape. In turn, adaptive approximations with MP algorithms are well known for their greediness (Baumgartner et al., 2013; Wacker and Witte, 2013). Another significant limitation of the time-frequency analysis of brain signals is its high computational complexity and the difficulty in interpreting the results by biologists and physicians. As a result, most of these applications are usually limited to scientific research. Therefore, for clinical applications, we still need to look for methods of exploring the single-trial evoked brain activity, which would utilise time–frequency analyses while avoiding high computational burden.

## 2. Aim

Since early processing of external stimuli is disturbed in schizophrenia and it plays an important role in the development of productive symptoms, obtaining as much information as possible about the type of these abnormalities is crucial for a good understanding of the pathology of schizophrenia. In our research on the evoked brain potentials registered from chronic schizophrenic patients, we observed changes in the high-frequency spectrum in comparison to healthy subjects (Maciejowski et al., 1987; Maciejowski, 1991; Małyszczak et al., 2003).

The aim of this study was to demonstrate differences in the spectral structure of single-trial visual evoked potentials (VEP) in patients with chronic schizophrenia in comparison to physiological recordings, thus confirming the presence of neuronal distraction in the course of chronic schizophrenia. The second aim was to show how to describe these abnormalities by applying relatively simple methods of signal processing, the tools of data visualization in the time and frequency domain, and statistical analysis.

## 3. Material

We used archival electroencephalographic data registered in the Laboratory of Evoked Potentials at the Department of Pathophysiology, Wrocław Medical University with the use of a specially designed system for measuring the evoked activity of the human brain, i.e., STimulated ELectroencephalogram on-Line Analyser (STELLA) (Maciejowski, 1986; Jagielski and Maciejowski, 1991). The system is based on a PC microcomputer, two 10-bit A/C converters, a multichannel biological enhancer with 100-Ohm input impedance, and the visual stimulator with a screen on which we presented pattern-reversal stimuli with black-and-white checkerboard pattern. The measurement was synchronized with the moment of transferring the checkerboard pattern from positive to negative. We used a square pattern of 2 cm that covered the whole stimulator screen. Patients kept their eyes on a tag placed in the middle of the screen. Pattern reversal stimulation is a technique commonly used to stimulate VEP in research on schizophrenia (Butler and Javitt, 2005; Rady et al., 2011; Babhulkar et al., 2017). We applied the visual stimulation without the commonly used oddball paradigm, to obtain unattended VEP, representing early signal processing. The measurement was made in a bipolar system on both hemispheres with the use of Cz-O1 and Cz-O2 leads and with the ground electrode placed on a patient’s forehead. These parieto-occipital leads are appropriate for observing the gamma-band of visual responses in the brain (Gruber et al., 1999). We used coated silver-silver chloride cup electrodes, filled with conducting paste, and secured to the skin with a tape. At the time of recording, inter-electrode impedance was of approximately 5 Ohm. The sampling rate for each channel was 1 kHz, the single-trial time window of analysis was of 500 ms length, and the frequency of stimulation was equal to 0.5 Hz. An analogue network band-stop filter was used to remove power grid (50 Hz) artefacts from the measured signals. At the time of measurement, we detected measurement artefacts online with the use of a threshold discriminator that rejected responses of amplitudes exceeding the linear activity of the biological enhancer. The rejection of ten subsequent measurements resulted in the automatic cessation of the examination. Examinations were conducted in a darkened room and the examined patients were recorded with a camera and monitored on the screen. If patients closed their eyes or moved their heads, which was interpreted as lack of cooperation with the examiner, the recording was stopped. Each technically correct visual cortex response to stimulation was recorded directly on the computer’s hard drive. We registered 128–300 single-trial visual responses during each recording session.

The VEP signals were collected from patients with schizophrenia and from healthy individuals. We chose the group of 19 adults (15 females, 4 males), age 22–50 years (median 30), diagnosed with paranoid schizophrenia according to DSM-III-R and a psychiatrist. All of them experienced visual hallucinations in the past, therefore this group is representative for studying visual-information early-processing disorders in schizophrenia. The subjects were not under the influence of psychotropic drugs, all were free of any metabolic, neurological or ophthalmic disease which might affect the results of the experiment, and there were no neurological problems or head injuries in their medical interview. The control group consists of 19 healthy adults (11 females, 8 males), age 22–50 years (median 33). All examined subjects were right-handed.

## 4. Methods

No post-measurement artefact corrections and other pre-processing methods were performed on the measured data sets, to maintain the original information of the individual signals.

Initially, the evoked responses were averaged to perform a classical electrophysiological evaluation of VEPs, which was performed by a neurophysiologist with many years of experience in describing this type of signals. Next, single-trial potentials were analysed. The evolution of single-trial brain responses was examined by graphically representing non-averaged trials in the time domain, using colour images comprising lines, with signal amplitude in each line matched by the colour scale (Fig. 1). Individual trials were arranged in the chronological order (from the first to the last stimulation) or ranked according to the specified criterion (e.g., the stage or the latency of the maximal amplitude in a time period). For this purpose, we used the own-designed computer programme written in FORTRAN and introduced by Maciejowski (1986). Such a visual representation of non-averaged VEP signals allows for the evaluation of changes that occur at the time of generating brain responses to a particular stimulus and of their repeatability.

**Figure 1:**
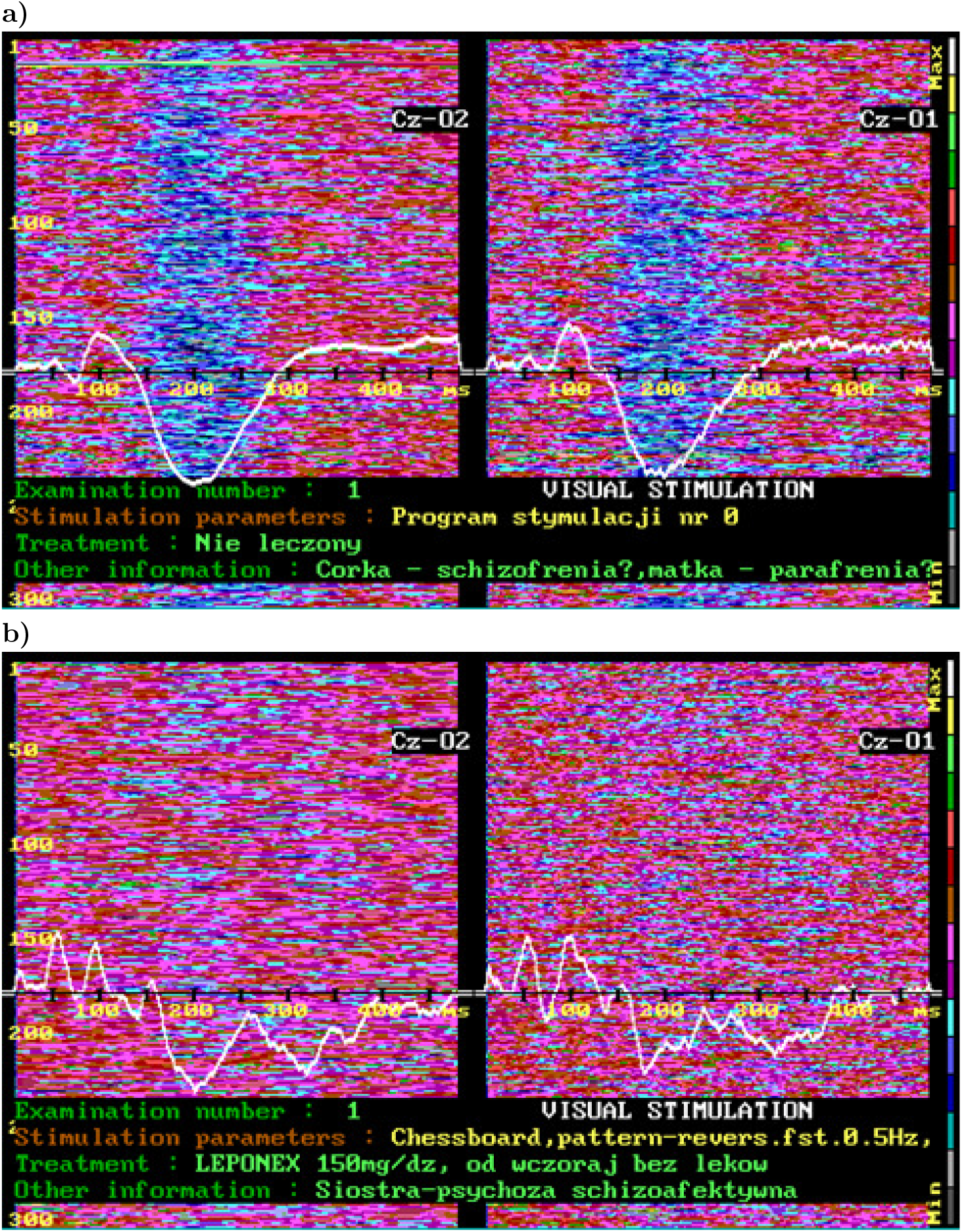
Single-trial brain evoked responses presented in the form of colourful pictures consisting of lines, where colours (see colour bar on the right side) match signal amplitudes. For healthy individuals (a) the synchronization of the response to visual stimuli is significantly higher than in patients with schizophrenia (b). The white line presents the averaged VEP obtained from 300 single responses registered synchronously with pattern-reversal stimuli, and it is physiological in (a) and abnormal in (b). Screenshots from the STELLA measurement system.

In order to determine the frequency structure of the VEP signals we transformed the single-trial VEP from the time domain to the frequency domain by applying the fast Fourier transform (FFT) (Brigham and Oran, 1988; Burrus, 2012). Having records of 500-ms length we performed the FFT calculation separately for each single trial, estimating the spectral density with 2-Hz (1/500 ms) resolution. We analysed the power spectrum within 2–200-Hz frequency range, which involves all EEG characteristic frequencies, i.e., delta, theta, alpha, slow beta, fast beta, and gamma. For calculations, we used our own scripts written in MATLAB (MathWorks, USA) with the use of the Signal Processing Toolbox (MathWorks, USA). To visualize the results and to allow for the evaluation of the trial-to-trial variability of VEP power spectra as well as the hemispheric asymmetry, we plotted 3D spectrograms (see Fig. 2 and Fig. 3).

**Figure 2:**
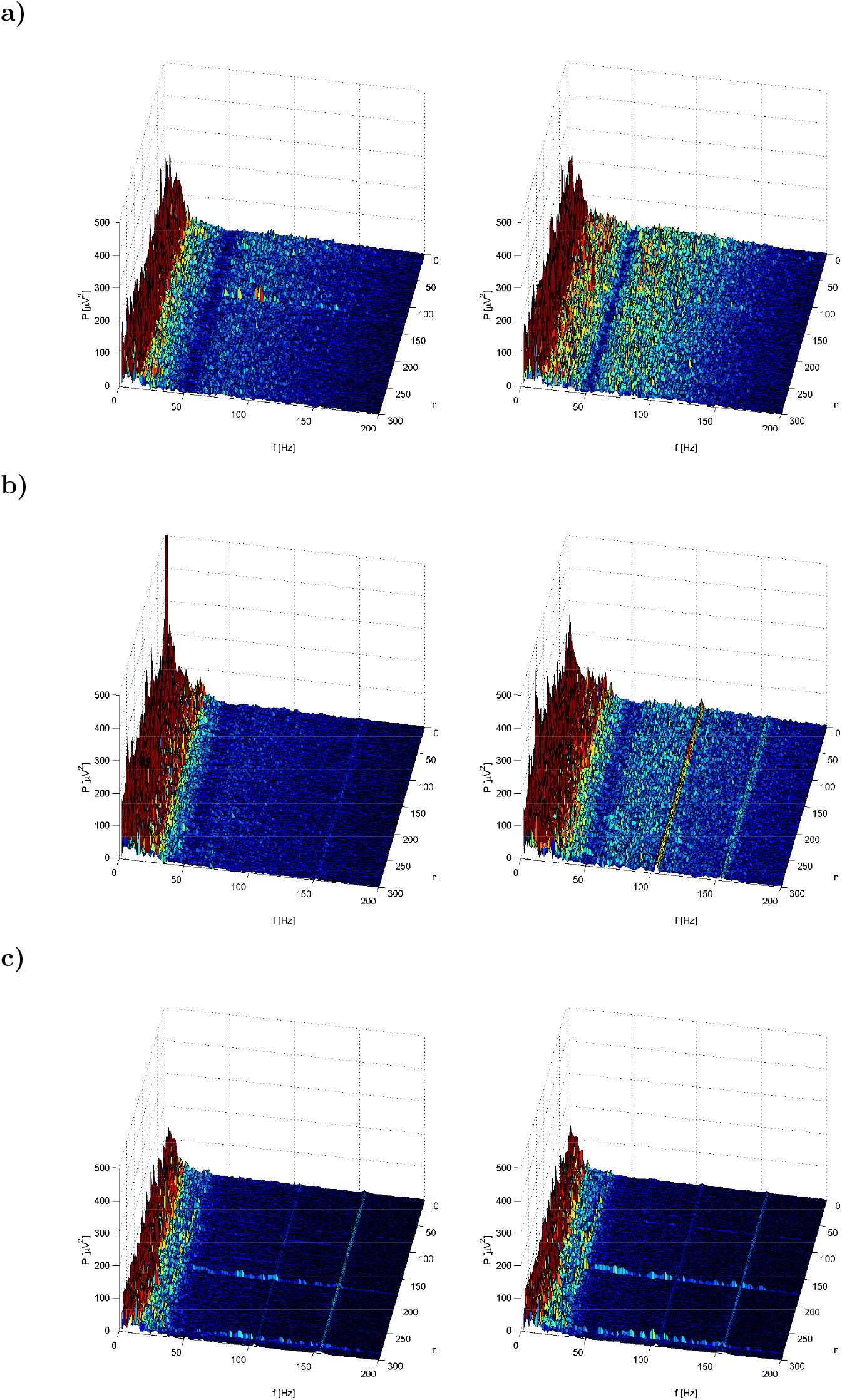
Power spectrograms of single-trial visual responses evoked in the human brain in healthy individuals at the time of examination (300 repetitions of the identical pattern-reversal stimuli). Index *n* denotes consecutive trials of 500-ms length. The left column corresponds to the left hemisphere and the right column to the right hemisphere. Consecutive rows correspond to selected individuals. The high power of the signal is visible in lower frequencies, which is an expected result, because the power spectrum of physiological EEG is similar to that of pink noise with its 1/*f* characteristics. In frequencies exceeding 35 Hz, power spectrum is very weak, except for the fundamental frequency of the power network (50 Hz—cut out with an analogue band-stop filter) and its harmonics. In (a) and (b), we can observe a small hemispheric asymmetry in the form of higher amplitudes at higher frequencies for the right (not dominating) hemisphere.

**Figure 3:**
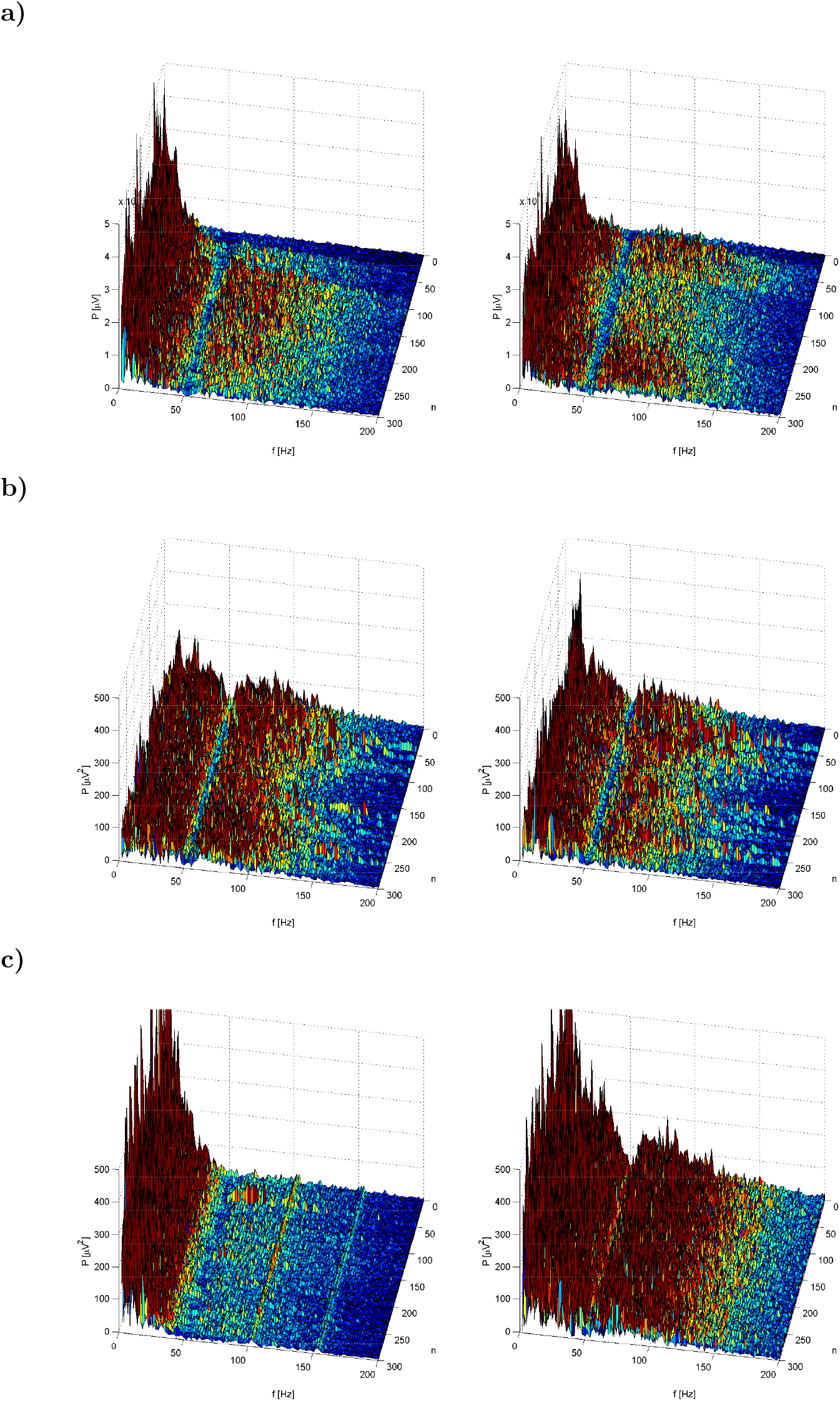
Power spectrum of single-trial VEPs from three different patients with schizophrenia. Index *n* denotes consecutive trials of 500-ms length. The left column corresponds to the left hemisphere and the right column to the right hemisphere. Consecutive rows correspond to selected individuals. In comparison to healthy individuals (Fig. 2), the power of response in high frequencies (> 36 Hz) is much larger in both hemispheres in (a) and (b), and in the non-dominating hemisphere of the patient from (c). The effect of the analogue band-stop filter in the form of weakening the signal at the power-line frequency of 50 Hz is clearly visible, though the two higher harmonics are noticeable.

Next, the obtained spectral power values for specific frequencies were subjected to quantitative analysis (QEEG) to compare the absolute and the relative power of a signal in the in the following spectral ranges: 2–36 Hz, 62–96 Hz, and 106–140 Hz. It should be noted that a routine EEG analysis involves only the first frequency band (2–36 Hz, containing the delta, theta, alpha, and beta frequency bands) of power spectrum, since rhythm patterns of basic brain bioelectric activity are of slow type, and EEG power spectrum is comparable to the power spectrum of pink noise. The next two frequency ranges (62–96 Hz and 106–140 Hz) were analysed to determine changes in the gamma band. We chose the above sub-bands to avoid distortions arising from the power supply network (50 Hz) and from its band-stop filtering on the results. It was also desirable to visibly separate the examined sub-bands. Frequency bands of the same width allowed for obtaining consolidated results for each band of identical weight. The relative spectral power was defined as the proportion of the power expressed in a given frequency band to the total spectral power of signal within the range of 2–200 Hz, i.e., as the percentage of energy transmitted in a specific frequency band to the total signal energy. Thus, we could statistically analyse the intensity of the signal within specific frequency bands and eliminate the effect of signal amplitude on results. Calculations were done for the sum of the Fourier coefficients of a given frequency band, summing them up separately for each patient and for each single-trial VEP. In case of the relative power spectrum, we calculated the ratio of the sum of Fourier coefficients from a specific frequency band and the sum of coefficients from the entire 2–200-Hz power spectrum. Using statistical analysis, we compared the results obtained for the group of patients with schizophrenia with those from healthy individuals, separately for each frequency band and EEG lead (the latter being related to either the left or the right hemisphere). We used the two-sample dependent *t*-test for equality of means, which was preceded by the Kolmogorov–Smirnov test and the Lilliefors test for normality, as well as by the Levene’s test to assess the equality of variances. In all tests, a significance level *α* = 0.05 was applied. The statistical analysis was done in STATISTICA (StatSoft, v. 10) (Nisbet et al., 2009).

## 5. Results

When analysing trial-averaged VEPs in schizophrenic patients, no P100 amplitude reduction was observed, and prolonged latencies were frequently detected for the N200 waveform only, similarly like in Małyszczak et al. (2003). However, morphological changes were noticed in patients with schizophrenia. A good example is presented in Fig. 1b, where the averaged VEPs show the presence of the P100 and N200 components with correct amplitude and latency, however, the entire waveform is rather chaotic, with several large amplitude changes over time. Such abnormal VEP morphologies do not occur in healthy subjects and have been observed in schizophrenia (Knebel et al., 2011; Babhulkar et al., 2017).

The trial-to-trial variability of brain evoked potentials registered directly after visual stimulation has been assessed qualitatively using their graphical representation, like in. Fig. 1. This allows for monitoring the level of instability of single responses to the repeated stimulus in normal and pathological cases. In healthy individuals (Fig. 1a)), we observe a relative time-stability of recorded VEPs with respect to the onset of the stimulus, resulting in undisturbed development of particular components of the trial-averaged evoked response, e.g., the P100/N200 waveform. This stability indicates physiological repeatability of singletrial evoked responses, resulting from synchronized processing of the same stimulus. The picture obtained for a patient with schizophrenia (Fig. 1 b)) demonstrates wide variations in single-trial VEPs, which, after averaging, resulted in a trace of clearly different morphology. Through the qualitative analysis of such figures we conclude that the synchronization of neural responses to visual stimulus is significantly better in healthy individuals than in patients with schizophrenia.

After transforming single-trials into the frequency domain using FFT, spectrograms were computed, presenting the obtained spectra in the domain of subsequent trials. Illustrative results are presented in Fig. 2 (healthy individuals) and Fig. 3 (patients with schizophrenia), where consecutive rows correspond to selected individuals. The occupied frequency range was rather narrow in the group of healthy individuals, i.e., almost the entire signal energy was concentrated <36 Hz. The low percentage of high frequencies for physiological VEPs is a natural phenomenon, following the distribution of the physiological EEG spectrum, which resembles pink noise (Fig. 2) with its 1/*f* characteristics. As opposed to healthy individuals, patients with schizophrenia demonstrate considerable energy in high frequencies, i.e., large amplitudes of the spectral power are visible for gamma frequencies, and the spectra fade away slowly around 150 Hz. This can be observed bilaterally (Fig. 3a and b) or with hemispheric asymmetry (Fig. 3c)).

Analysing the absolute spectral power in the 2–36-Hz band, there are no statistically significant differences between the schizophrenic and the control group in the left (dominant for all subjects) hemisphere, whereas in the right, non-dominant hemisphere, the spectral power in healthy subjects is statistically significantly higher than in schizophrenic patients (first column from Fig. 4). This pattern occurs in the commonly analysed range of the EEG spectrum, including the delta, theta, alpha, and beta bands. For higher frequencies, including the gamma band, i.e., 62–96 Hz and 106–140 Hz, the absolute spectral power in the dominant hemisphere (Fig. 4a) is statistically significantly higher in schizophrenic patients than in healthy subjects. In the non-dominant hemisphere, the mean power spectrum in healthy subjects is larger compared to that in patients with schizophrenia, however, this difference is statistically significant in the 62–96-Hz range, yet, not statistically significant in the 106–140-Hz range (Fig. 4b).

**Figure 4:**
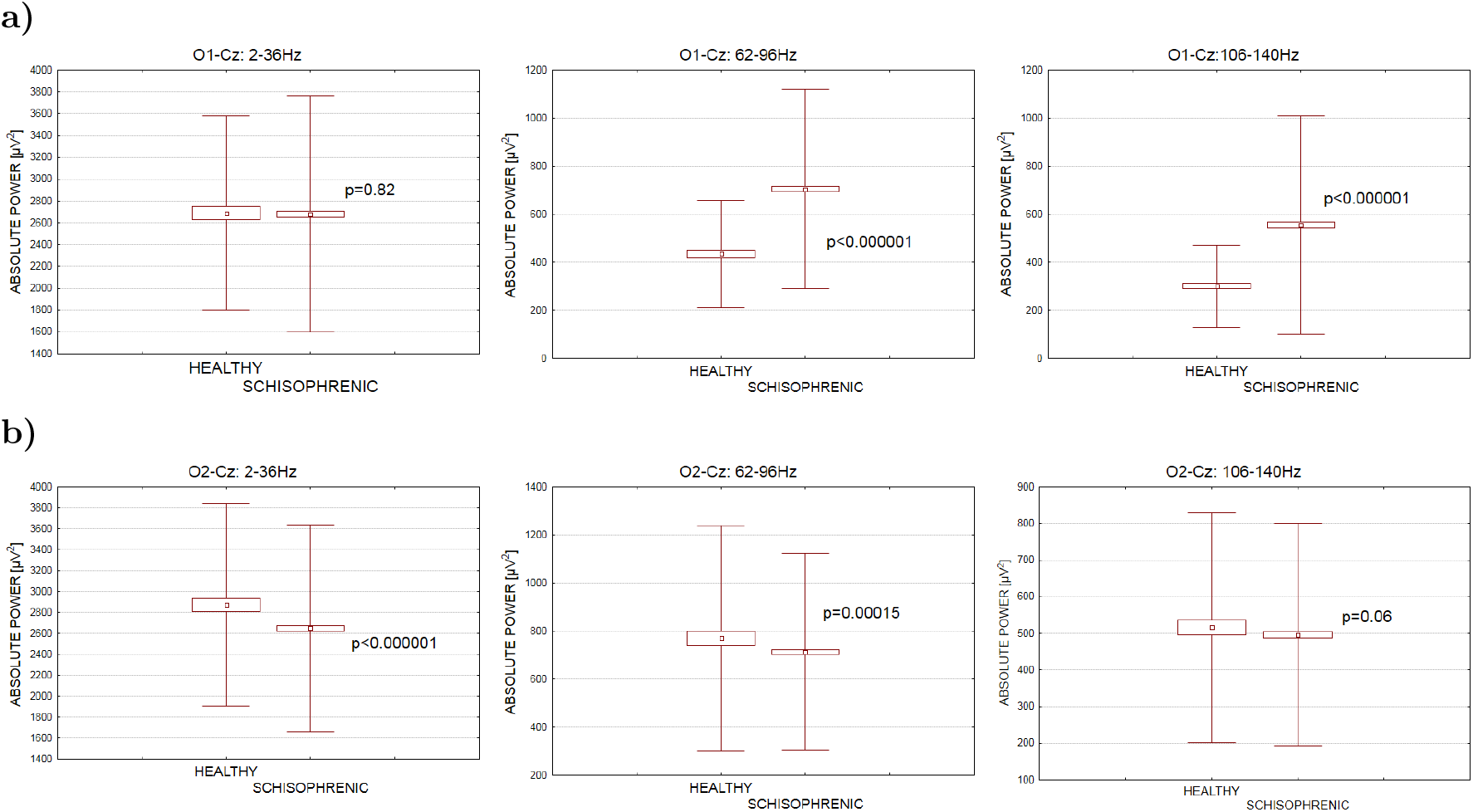
A comparison of statistical parameters for the absolute power spectrum of single-trial VEPs [*μ*V^2^] in the group of patients with chronic schizophrenia compared to healthy individuals. a) right- and b) left-hemisphere leads. The calculated mean values, confidence intervals, and standard deviations with *p*-values from *t*-test are given.

Analysing the relative spectral power (Fig. 5), the frequencies in the delta, theta, alpha, and beta bands (2–36 Hz) dominate in healthy subjects on both sides and these are statistically significant differences. However, for both the dominant (Fig. 5a) and the non-dominant hemisphere (Fig. 5b), in both high-frequency ranges (62–96 Hz and 106–140 Hz), the relative spectral power is statistically significantly greater in the chronic schizophrenia group compared to the control group. Thus, both absolute and relative spectral power in high frequencies is significantly greater in the dominant hemisphere in schizophrenics. For the non-dominant hemisphere, this is not visible when we analyse the absolute spectral power, whereas when analysing the relative spectral power, statistically significant differences of this kind were detected in both considered high-frequency bands.

**Figure 5:**
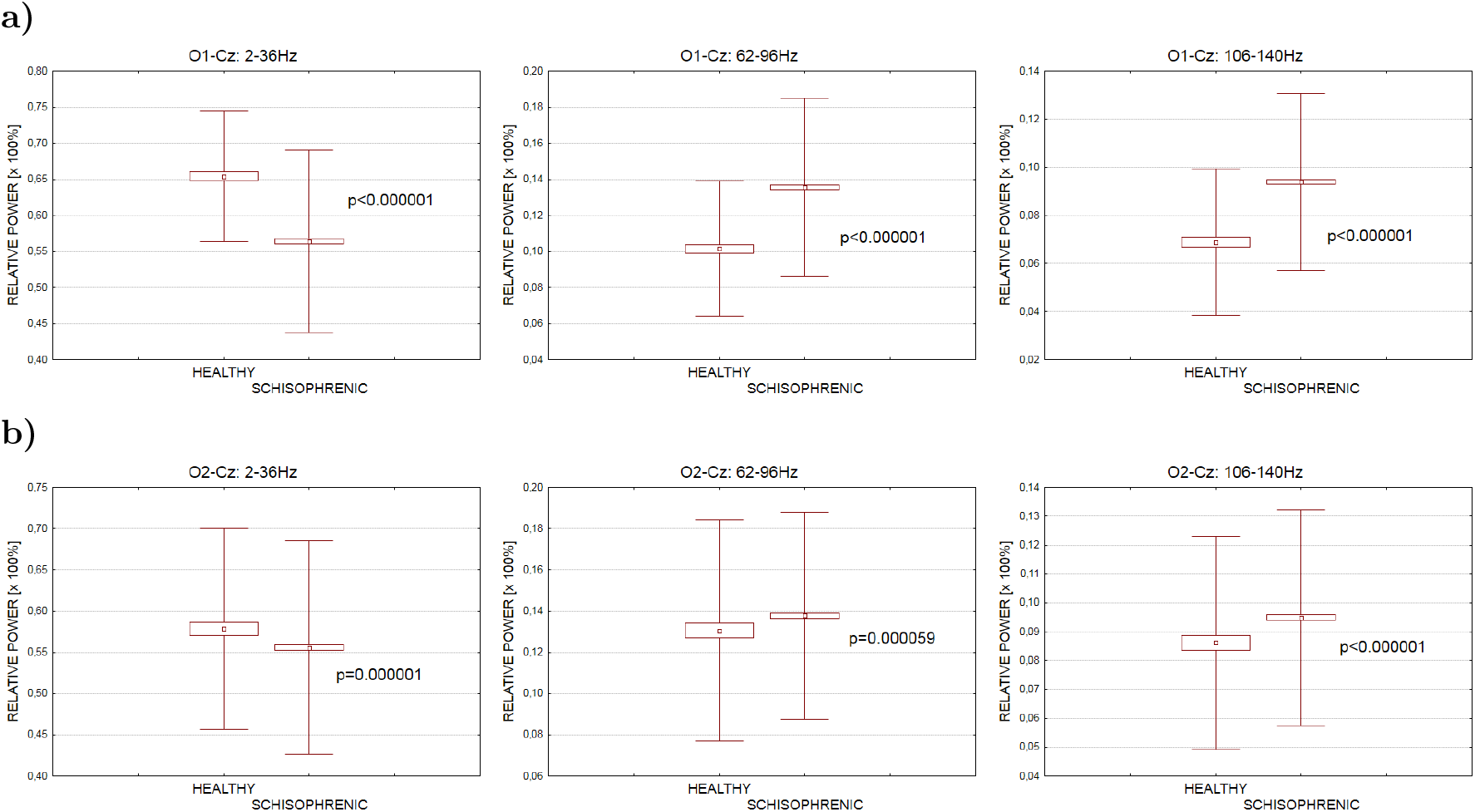
A comparison of statistical parameters for the relative power spectrum of single-trial VEPs [%] in the group of patients with chronic schizophrenia compared to healthy individuals. a) right- and b) left-hemisphere leads. The calculated mean values, confidence intervals, and standard deviations with *p*-values from *t*-test are given.

## 6. Discussion

In recent decades, a number of neurophysiological abnormalities in patients with schizophrenia have been identified and intensively studied. The most studied were: abnormalities in smooth pursuit eye movements, P50 sensory gating, prepulse inhibition, P300 amplitude and latency, mismatch negativity, and neural synchrony. They were characterized by low dependence on the severity of the psychotic state, and as more frequent in relatives of sick people. They have been investigated as potential biomarkers or endophenotypes of schizophrenia (Onitsuka et al., 2013).

Many studies report impaired processing of the visual stimuli in patients with schizophrenia, which is confirmed by measurements of evoked potentials. This dysfunction results from disorganized neural network responsible for experiencing sensual impressions, which can explain some symptoms in schizophrenia (see Section 1.2, and references therein). Especially, the changes in gamma oscillations are recently studied, due to their role in neuronal synchronization in attention and memory, and to their association with brain neurotransmitter systems relevant for schizophrenia (Engel et al., 2001; Lee et al., 2003; van der Stelt et al., 2004; Light et al., 2006; Başar-Eroglu et al., 2007; Womelsdorf et al., 2006; Fries et al., 2007; Spencer et al., 2008; Uhlhaas and Singer, 2010; Andreou et al., 2014; Hirano et al., 2015; McNally and McCarley, 2016; Klein et al., 2020; Sauer et al., 2020; Tada et al., 2020). Interestingly, the spontaneous gamma activity may be increased in schizophrenia, reflecting a disruption in the normal balance of excitation and inhibition, and can interact with evoked oscillations, possibly contributing to their deficits found in gamma-band in chronic schizophrenic patients (Hirano et al., 2015). Multiple task-related studies in patients with schizophrenia show reduced signal power in gamma frequency band and reduced gamma synchronization. Andreou et al. (2014) suggest that this does not apply to the resting EEG, and the increase in connectivity in the gamma band is not non-specific, but involved in a strongly lateralized network covering areas of the cerebral cortex essential for language and memory functions, especially impaired in patients with schizophrenia. These authors suggest that resting-state studies might reveal new aspects in the pathophysiology of schizophrenia (Andreou et al., 2014). Moran and Hong in their summarized paper suggested that the gamma oscillation is often the first component of the response to a sensory stimuli (Moran and Hong, 2011).

To gain a better understanding of these phenomena, we have analysed the high-frequency (gamma) changes in single-trial (non-averaged) visual evoked potentials collected from the group of chronic schizophrenic patients with visual symptoms and from healthy individuals. Similarly to many other authors we used parieto-occipital leads from both hemispheres for comparison. We chose the spectral analysis based on the short-time Fourier transform in place of more complex techniques of time–frequency signal decomposition, and spectrograms as a visualization tool.

In contrast to other studies, where slowing down of brain activity was found in spontaneous EEG in schizophrenia (see Section 1.1 and references therein), we observed an increased spectral power in high frequencies, i.e., 62–96 Hz and 106–140 Hz. This observation is similar to the findings presented by Andreou et al. (2014) and Hirano et al. (2015) for auditory stimuli, yet, our experiments and methodology were different. Although our experiment, too, is based on a stimulus-dependent paradigm, we used a visual stimulus and obtained early stimulus-processing potentials, not the steady-state responses. On the other hand, single-trial VEPs contain considerable amounts of spontaneous EEG activity in the signal energy. Even though we found increased gamma power in the left-hemisphere lead only when the absolute spectral power was the measure, for the relative spectral power high frequencies in both analysed ranges of gamma frequencies revealed significantly greater power bilaterally in schizophrenic patients.

It is difficult to say to which extent the power-spectrum findings for the two high-frequency bands are influenced by the stimulation, since the frequency profile of a typical VEP is concentrated at relatively slow oscillations (see white traces in Fig. 1). Therefore, it is possible that the observed differences between the patients and the healthy subjects reflect spontaneous or induced (Pfurtscheller and Lopes da Silva, 1999; CatherineTallon-Baudry and Bertrand, 1999; Engel et al., 2001) rather than evoked activity. In our study, we observed higher average spectral power of single-trial VEPs in patients with schizophrenia, but it was not related to the higher amplitude of the average waveforms (see Fig. 4 and Fig. 5 vs Fig. Fig. 1). However, visual evoked responses are more disorganized and less repeatable in schizophrenia than in healthy individuals (Knebel et al., 2011). In our study, single responses in patients with schizophrenia were more intense (amplitudes were higher) but, after averaging, they resulted in no greater VEPs than in healthy individuals. This paradox can be explained by the fact that single responses in patients with schizophrenia are of greater diversity, which, after averaging of results and in comparison to healthy individuals, show flattened shape, which indicates a greater participation of noise with zero expected value and a limited participation of repeatable components with non-zero expected value. It results directly from the properties of the averaging method (König et al., 2015). Moreover, it is worth noting that research on auditory evoked potentials showed impairment of habituation in patients with schizophrenia (Moran et al., 2012), and the habituation phenomenon could have an impact on the result too. Finally, we must point out that when the Fourier transform is applied, it is tacitly assumed that single-trial VEPs are stationary. However, this assumption may not be met even for encephalographic signals as short as of 500-ms length and so the nonstationarity may have biased our results somewhat (Kipiński et al., 2011; Kipiński and Kordecki, 2021).

When comparing our results with those from other studies it is important to point out that even though we observed changes in high (gamma) frequencies, we analysed power spectra rather than gamma oscillations as such. It would be worth comparing the results of our observations with the changes in state (amplitude or phase) of the high frequency oscillations and with the resting-state EEG from the same group of patients. Unfortunately, our measurements did not allow such observations because in the collected signals only short fragments of the EEG immediately following the stimulus presentation were recorded, without the prestimulus or resting-state period.

Gamma oscillations are related to attention and memory, but they are also connected to the function of early visual cortex (Klein et al., 2020) and thalamo-occipital cortices (Sauer et al., 2020). Besides, spinothalamic tract is believed to be the place where VEPs for unattended stimuli are generated (Lalor et al., 2012). Thus, the occurrence of high-frequency changes in single-trial VEPs for unattended stimuli entitles to pose the question why high frequencies are excited, since in other studies on schizophrenia impaired gamma oscillations were found. We think that this result may be influenced by a high content of induced or spontaneous activity in single trials and the nonstationarity.

The presented study broadens our understanding of high EEG frequencies in chronic schizophrenia and may constitute a starting point for further comparative studies. It would also be worth checking whether proposing a parametric modelling of the spectrum trial-to-trial changes of single-trial VEP, as proposed by Kipiński and Kordecki (2021), would provide new information on the behaviour of high-frequency oscillations in patients with schizophrenia.

## Acknowledgements

We dedicate this work to the memory of Professor Józef Jagielski (died on March 13, 2021), a long-term head of the Department of Pathophysiology at the Wrocław Medical University, a pioneer in the study of brain evoked potentials.

